# Benchmarking Automated Cell Type Annotation Tools for Single-cell ATAC-seq Data

**DOI:** 10.1101/2022.10.05.511014

**Authors:** Yuge Wang, Xingzhi Sun, Hongyu Zhao

## Abstract

As single-cell chromatin accessibility profiling methods advance, scATAC-seq has become ever more important in the study of candidate regulatory genomic regions and their roles underlying developmental, evolutionary and disease processes. At the same time, cell type annotation is critical in understanding the cellular composition of complex tissues and identifying potential novel cell types. However, most existing methods that can perform automated cell type annotation are designed to transfer labels from an annotated scRNA-seq data set to another scRNA-seq data set, and it is not clear whether these methods are adaptable to annotate scATAC-seq data. Several methods have been recently proposed for label transfer from scRNA-seq data to scATAC-seq data, but there is a lack of benchmarking study on the performance of these methods. Here, we evaluated the performance of five scATAC-seq annotation methods on both their classification accuracy and scalability using publicly available single-cell datasets from mouse and human tissues including brain, lung, kidney, PBMC and BMMC. Using the BMMC data as basis, we further investigated the performance of these methods across different data sizes, mislabeling rates, sequencing depths and the number of cell types unique to scATAC-seq. Bridge integration, which is the only method that requires additional multimodal data and does not need gene activity calculation, was overall the best method and robust to changes in data size, mislabeling rate and sequencing depth. Conos was the most time and memory efficient method but performed the worst in terms of prediction accuracy. scJoint tended to assign cells to similar cell types and performed relatively poorly for complex datasets with deep annotations but performed better for datasets only with major label annotations. The performance of scGCN and Seurat v3 was moderate, but scGCN was the most time-consuming method and had the most similar performance to random classifiers for cell types unique to scATAC-seq.

## 1 Introduction

With the advancement of single-cell sequencing technologies, researchers not only can profile single-cell transcriptomes by scRNA-seq, but can also measure multiple modalities at the single-cell level (Packer and Trapnell, 2018; Carter and Zhao, 2021), among which scATAC-seq is probably the most widely used sequencing technology (Buenrostro et al., 2015; Cusanovich et al., 2015). scATAC-seq can quantify chromatin accessibility across tens of thousands of single cells and is an important tool to study gene regulation accompanied with scRNA-seq (Buenrostro et al., 2018; Fiers et al., 2018; Jia et al., 2018; Wang et al., 2022). Single-cell studies usually start with cell type annotations and accurate and robust annotations are crucial for downstream functional analyses that are often conducted in a cell-type-specific manner. Cell type annotation is often laborious and involves automated annotations from computational tools followed by verification and manual annotations from experts (Clarke et al., 2021). Although there are many tools designed for automated cell type annotations for scRNA-seq data (Abdelaal et al., 2019; Pasquini et al., 2021), only a limited number of tools are available and suitable for scATAC-seq data. As scATAC-seq becomes more mature and widely adopted in single-cell studies, there is a need to comprehensively evaluate their performance on annotating scATAC-seq data.

Currently, there are two types of annotation tools that can be applied to scATAC-seq data. The first category includes those originally designed for scRNA-seq data (intra-modality annotation), such as Seurat v3 (Stuart et al., 2019), Conos (Barkas et al., 2019) and scGCN (Song et al., 2021). The second category includes tools designed specifically for scATAC-seq data or for cross-modality annotation. The two representative methods in the second category are scJoint (Lin et al., 2022) and Bridge integration (Hao et al., 2022). Unlike the other methods that directly transfer labels from scRNA-seq to scATAC-seq after unifying the feature set through gene activity calculation, Bridge integration leverages a multimodal data as a bridge, avoiding potential loss of information and incorrectness of assumptions on feature relationships when calculating gene activities.

In this study, we benchmark these scATAC-seq annotation tools using real single-cell datasets from various tissues with available cell type annotations as the ground truth. The real data we considered included both paired data where scATAC-seq and scRNA-seq were simultaneously measured in each single cell and unpaired data where scATAC-seq and scRNA-seq were separately measured from the same tissue. We evaluated the performance of different methods on both annotation accuracy and scalability. For accuracy, we considered both the overall accuracy as well as accuracy on ATAC-specific cell types. For scalability, we compared running time and memory usage across different datasets. Apart from evaluating real data across different tissues, we also investigated the model performance across different cell numbers, mislabeling proportions, sequencing depths and number of unique cell types using a well-annotated human bone marrow mononuclear cell (BMMC) multimodal data (Luecken et al., 2021). The results of our study offer a basis for future methodology development and provide a reference for users to choose appropriate tools for cell type annotation from scATAC-seq data.

## 2 Results

### 2.1 Performance across Different Tissues

In this study, we used data from five different tissues, including mouse lung (Consortium, 2018; Cusanovich et al., 2018), mouse brain (Chen et al., 2019; Ma et al., 2020), mouse kidney (Cao et al., 2018; Miao et al., 2021), human peripheral blood mononuclear cell (PBMC) (Granja et al., 2019) and human bone marrow mononuclear cells (BMMC) (Luecken et al., 2021) to benchmark five methods for automated scATAC-seq label annotation, including Conos, Seurat v3, scGCN, scJoint and Bridge integration. For mouse lung, scRNA-seq data from both 10x Chromium (droplet-based) and Smart-seq2 (FACS-based) were collected. Among all the methods, only Bridge integration required multimodal data where scATAC-seq and scRNA-seq were simultaneously measured. Therefore, we collected multimodal data for each tissue except for mouse lung (the SHARE-seq data for mouse lung were sequenced too shallowly to be used). For the mouse brain, both SHARE-seq and SNARE-seq data were used as the multimodal data to benchmark Bridge integration separately.

We calculated three accuracy-related metrics on all ATAC cells, namely overall accuracy, weighted accuracy and F1 (macro) of precision and recall. For the first and third metrics, they were calculated based on predicted label, which was the cell type whose predicted probability was the largest. For weighted accuracy, we considered the similarity among cell types by calculating the weighted average of the entire predicted probability vector of each cell. Therefore, even though a predicted label was false, the score could be high if similar cell types had higher predicted probabilities. As can be seen from Figure 1A, all the five methods had better performance than plain K nearest neighbor (KNN) and the random classifiers. For mouse lung (both FACS and droplet) and mouse brain, scJoint had consistent and leading performance across all the three metrics, with only slightly lower F1 (macro) than scGCN on mouse brain. For the two human tissues (PBMC and BMMC), Bridge integration achieved the highest overall accuracy and F1 (macro); while for weighted accuracy, Bridge integration was the second best performer, following scJoint. For mouse kidney, there was no leading method across all three metrics, but scGCN and Seurat v3 had overall better performance.

**Figure 1.**
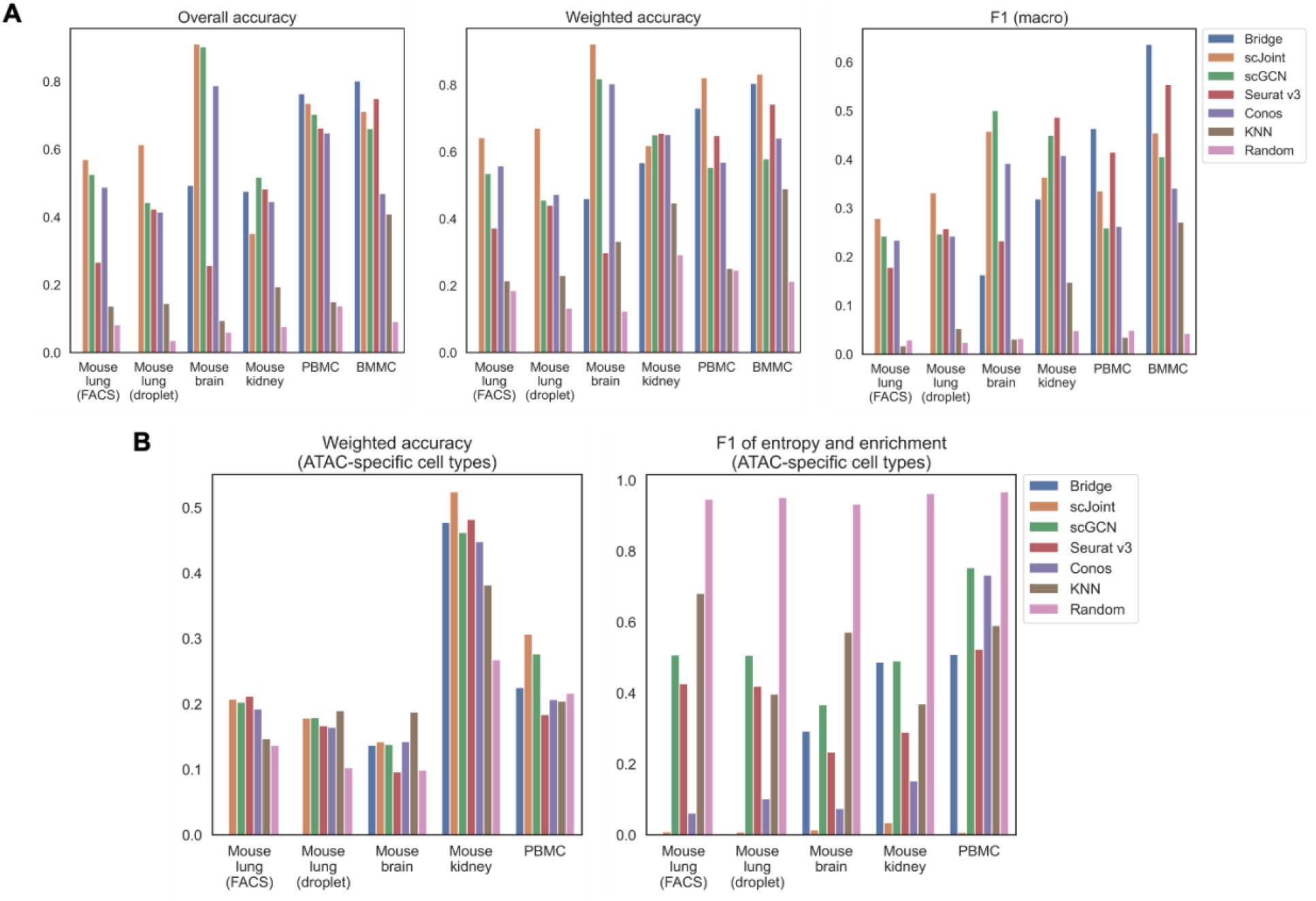
Performance of label transfer methods on single-cell data from selected mouse and human tissues. (A) Overall metrics considering performance on all scATAC-seq cells. (B) Metrics calculated on scATAC-seq cells labeled with ATAC-specific cell types. The Bridge results shown here for the mouse brain used SNARE-seq as the multimodal ‘bridge’. Comparison of results using SNARE-seq and SHARE-seq can be found in Supplementary Figure S1. For mouse lung (both FACS and droplet), Bridge integration was not considered because of no available multimodal data.

Apart from the three metrics assessing all ATAC cells, we designed two additional metrics for cell types that uniquely existed in ATAC data, namely weighted accuracy and F1 of entropy and enrichment (details in Materials and Methods). For ATAC-specific cell types, they could never be correctly classified because their labels did not exist in the reference RNA data. Then, there are two expected patterns for the predicted probability vectors of these cells. One is having predicted probability vectors close to the background distribution of cell types, and the other is having higher predicted probabilities for similar cell types in the RNA data. Both can have their own benefits in real practice. For example, for the first case, one can perform another round of manual annotations on cells with predicted probabilities close to the background distribution; while for the second case, one can tell from the predicted probabilities which existing cell types are the closest to the unknown cell type, however, this might suffer from misclassifying novel cell types due to biological similarity to known cell types. F1 (entropy and enrichment) and weighted accuracy were calculated over cells with unique cell types in the ATAC data to cover the first and the second cases, respectively (Figure 1B). Although scJoint consistently had relatively high weighted accuracy across tissues, there were not significant differences in weighted accuracy among all the five methods. For F1 (entropy and enrichment), the scores of scJoint and Conos were extremely low, while scGCN achieved the best scores among the five methods, followed by Bridge integration and Seurat v3.

Apart from the prediction accuracy, we evaluated the efficiency and scalability of the five methods by recording their running time and peak memory usage on each tissue (Figure 2). scGCN was the most time-consuming method and took the largest memory on mouse kidney, where there were about 71,000 cells in total. Conos was the most time and memory efficient method and remained nearly constant as the data scale increased. For the remaining three methods (Bridge integration, scJoint and Seurat v3), their running time didn’t differ significantly, but Bridge integration consumed more memory than others.

**Figure 2.**
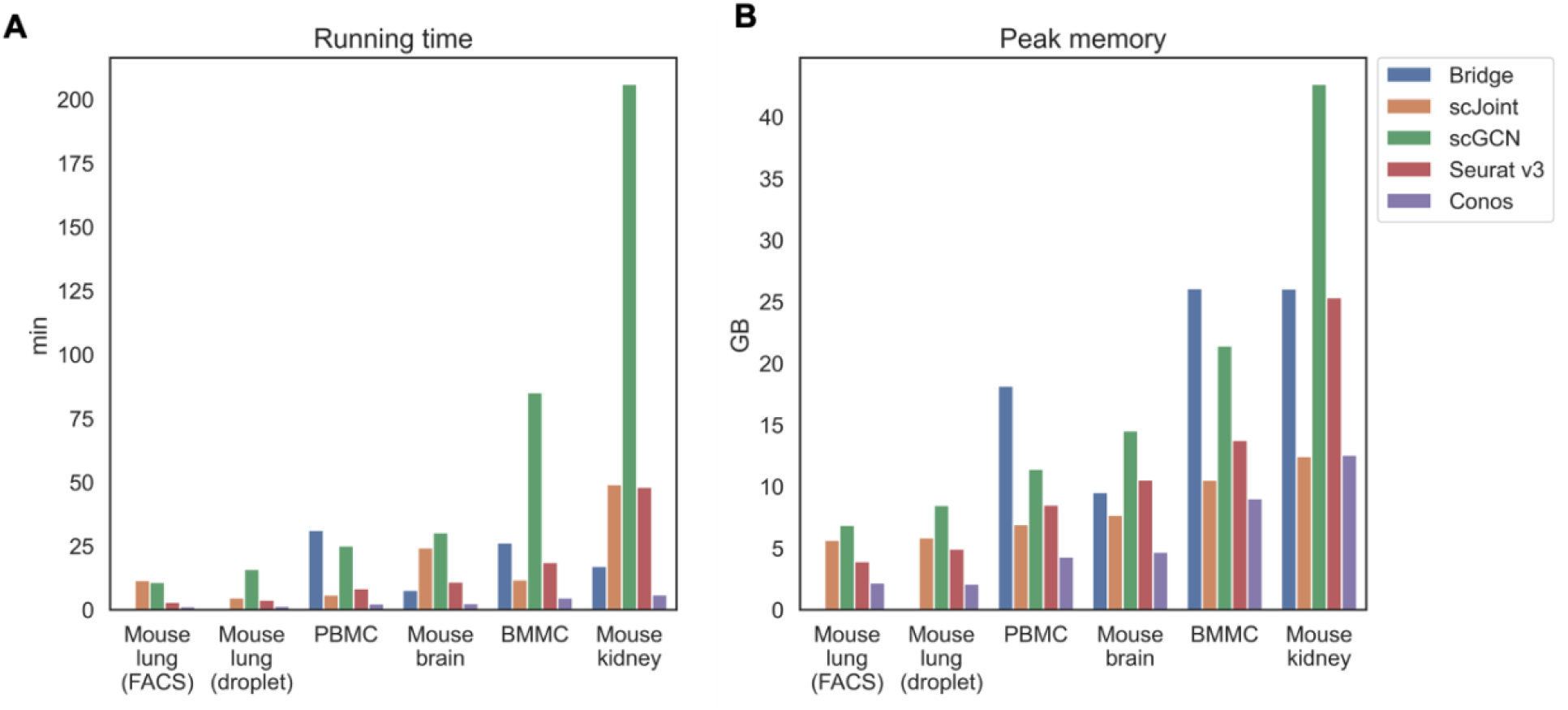
Running time (A) and peak memory usage (B) of different methods on selected tissues. Tissues are placed in the increasing order of their scales from left to right. For mouse lung (both FACS and droplet), Bridge was not considered because of no available multimodal data.

### 2.2 Performance across Different Data Scales

The BMMC data is a first-of-its-kind single-cell multimodal dataset which consists of about 70,000 cells with paired scRNA-seq and scATAC-seq measurements from 10 diverse donors at four sequencing sites. This dataset contains the largest number of cell types (22) among all selected tissues and captures both developmental and differentiated cell types. This dataset is the most comprehensive multi-modal benchmark dataset to date as far as we know, so we designed several experiments using the BMMC data to investigate the performance of different methods across diverse data characteristics. For all of the following experiments on the BMMC data, we manually separated all donors into three groups and used them as unimodal RNA data, unimodal ATAC data, and multimodal data, respectively (see Materials and Methods).

Figure 3A shows the performance of different methods across an increasing number of RNA cells, where KNN classifier and random classifiers were used as baseline references. We can observe that the value of three metrics didn’t further increase when the cell number reached 3k, which is a relatively small number given the current high-throughput sequencing technologies. In terms of overall accuracy and F1 (macro) of precision and recall, the order of the five methods from the best to the worst were the same, which was Bridge > Seurat v3 > scJoint > scGCN > Conos. For weighted accuracy, which took into consideration the similarity among cell types (see Materials and Methods for details) when assessing the predicted probability matrix, scJoint achieved the highest score and Conos was slightly better than scGCN, while the order of the rest of the methods remained the same.

**Figure 3.**
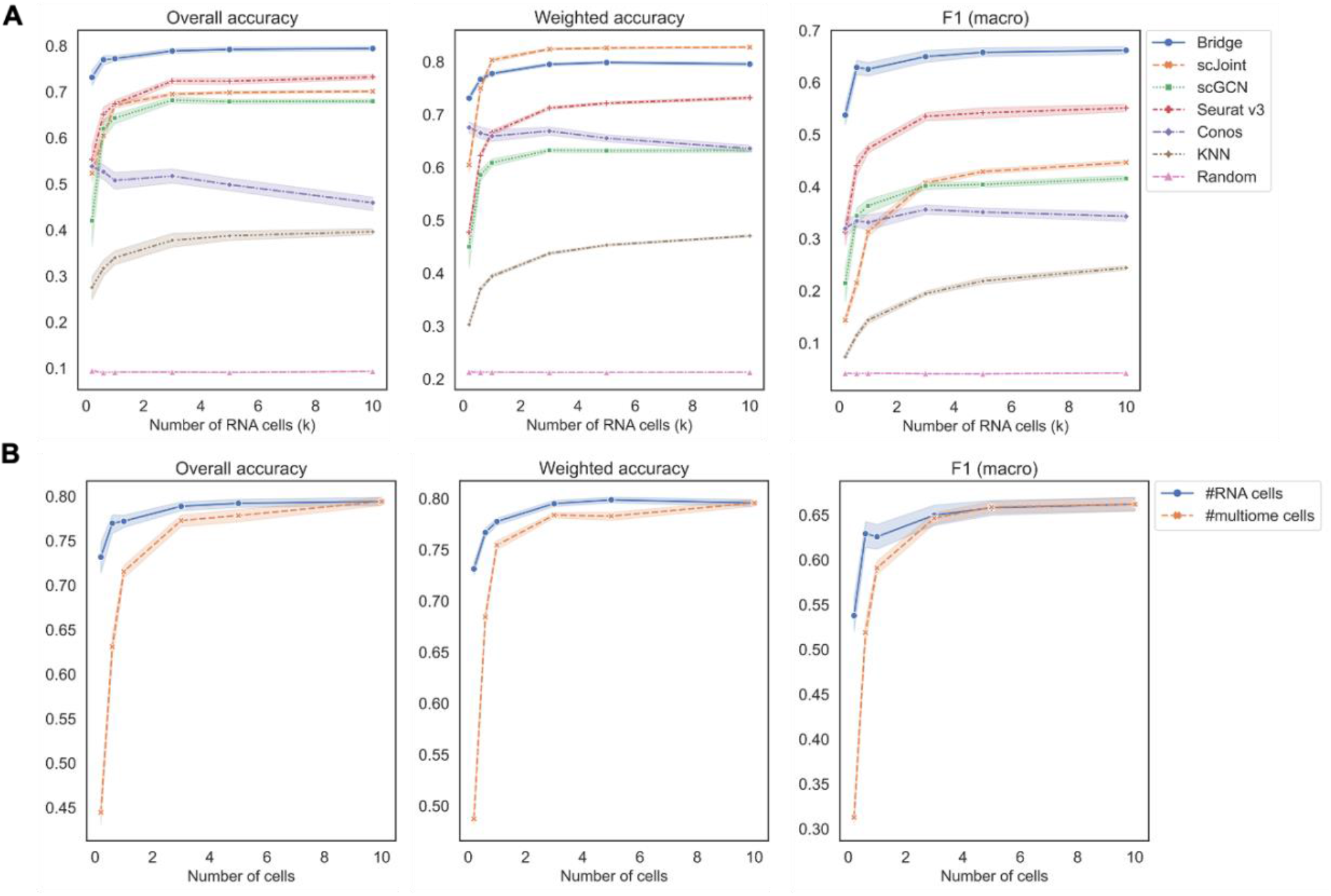
Performance of methods on different data scales of BMMC. (A) Change the number of cells in scRNA-seq only, while keeping scATAC-seq and multimodal (for Bridge only) cell numbers as 10k. Results shown here for Conos were paired with Pagoda2 for data processing. (B) Performance of Bridge while changing number of cells in scRNA-seq and in the multimodal data, respectively. The error band shows the 95% confidence interval.

Conos is a graph-based method and either Seurat or Pagoda2 is recommended for data processing before constructing the cell graph. We found the performance of Conos was worse when paired with Seurat (Supplementary Figure S2A), resulting in both lower values of the three metrics and higher instability. For Bridge integration, since it requires additional multimodal data as the ‘Bridge’, we performed another set of experiments specifically for Bridge by varying the number of cells in the multimodal data. We found the performance also stabilized when the cell number reached 3k and Bridge was more sensitive to the smaller number of cells in the multimodal data than in the unimodal RNA data (Figure 3B).

We recorded the running time and peak memory usage of the five methods when increasing the number of RNA cells (Supplementary Figure S3A). scGCN was the most time-consuming method and the second most memory-consuming method. Most of the time of running scGCN was spent on processing the data where intra-data and inter-data graphs were constructed. Bridge integration required the largest memory usage among all the methods because it involved additional multimodal data as the bridge, while its running time was close to that of scJoint. Conos and Seurat v3 were the two fastest methods and Conos was the least memory-consuming method.

### 2.3 Performance across Different Mislabeling proportions

The second set of experiments was designed to study the performance across different mislabeling proportions of the RNA data (Figure 4). For overall accuracy and F1 (macro), their scores remained constant for Bridge integration and Seurat v3 until mislabeling proportion reached 50% and decreased sharply when the proportion exceeded 70%. For scGCN, scJoint and Conos, their scores decreased slowly when the proportion was less than 50% and decreased faster after that. For weighted accuracy, almost all methods except scJoint decreased linearly as the mislabeling proportion increased. The order of the five methods was similar to the previous experiment, with Bridge and Seurat v3 being the top two methods in terms of overall accuracy and F1 (macro), and Conos and scGCN being the two worst-performers. scJoint was still the best method when considering the weighted accuracy. We also compared the performance of Conos when paired with Seurat and Pagoda2 separately and found that Conos (Seurat) was significantly worse than Conos (Pagoda2) across all metrics, especially when the mislabeling proportion was low (Supplementary Figure S2B).

**Figure 4.**
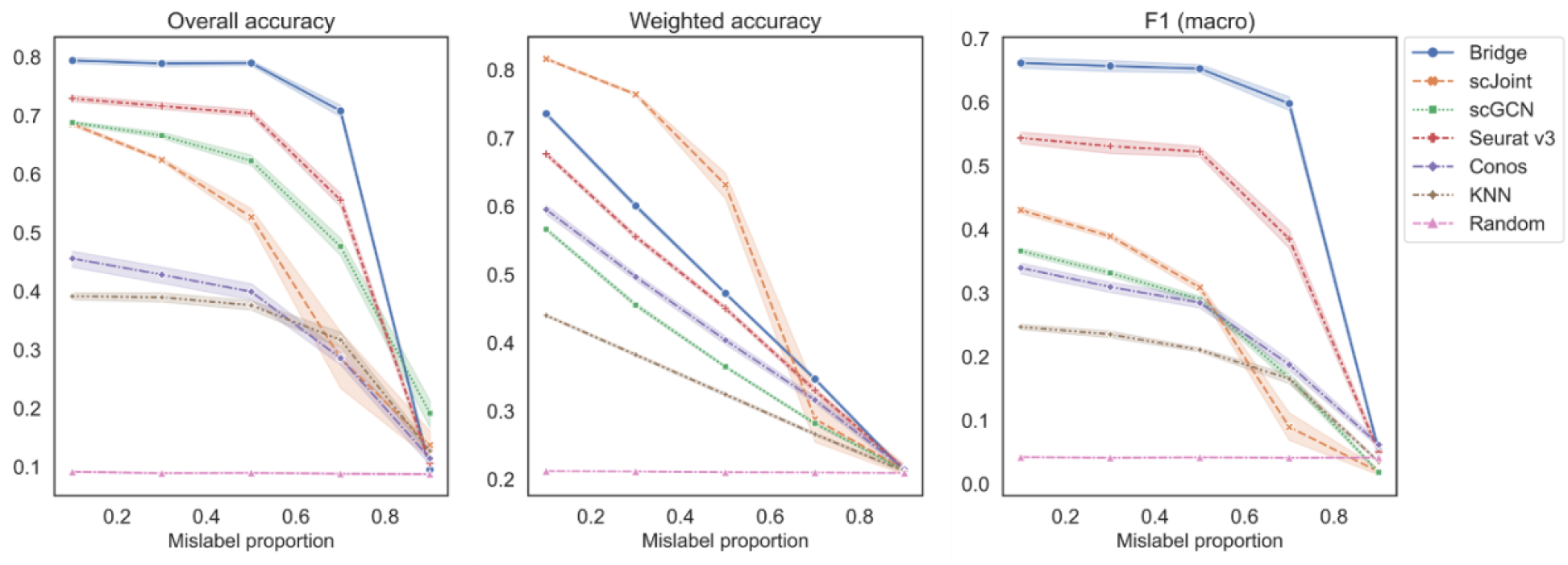
Performance of methods on different mislabeling proportions of BMMC. Results shown here for Conos were paired with Pagoda2 for data processing. The error band shows the 95% confidence interval.

### 2.4 Performance across Different Downsampling Proportions

In the third set of experiments, we downsampled each count matrix to some proportions to mimic different levels of sequencing depth. As we can observe from Figure 5A, all methods had a decreasing trend in their performance as the downsampling proportion decreased (lower sequencing depth). When the downsampling proportion was no less than 50%, the order of methods in terms of overall accuracy and F1 (macro) was the same, which was Bridge > Seurat v3 > scJoint > scGCN > Conos. When the sequencing depth was extremely low (downsampling proportion < 50%), Bridge integration was still the best performer, but the performance of scJoint became better than Seurat v3. As for weighted accuracy, scJoint was the best performer across all the methods. For Conos, its performance was worse when using Seurat for data processing compared to using Pagoda2 (Supplementary Figure S2C).

**Figure 5.**
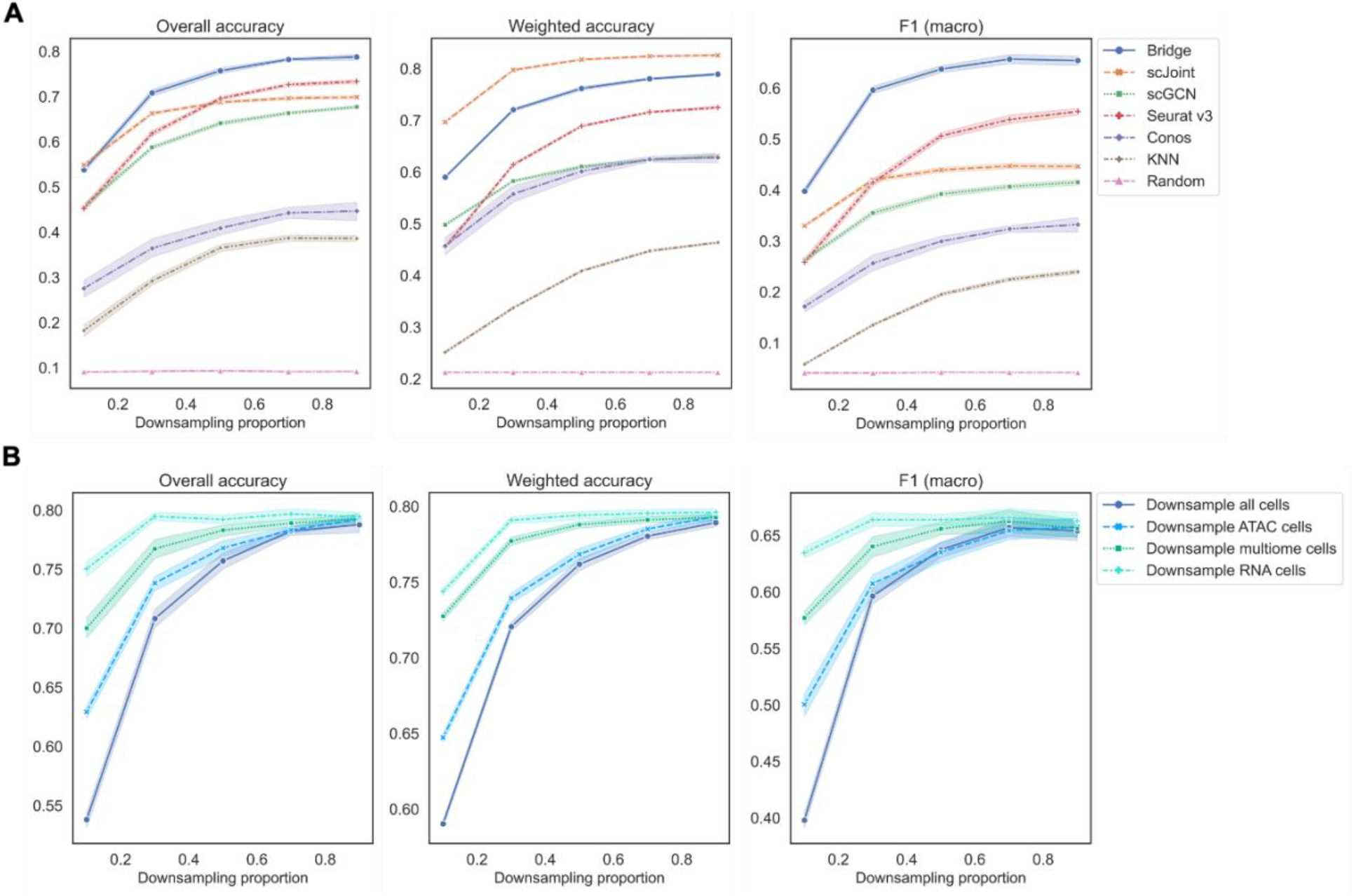
Performance of methods on different downsampling proportions of BMMC. (A) Downsample scRNA-seq, scATAC-seq and multimodal data (for Bridge only) at the same time. Results shown here for Conos were paired with Pagoda2 for data processing. Figures for scenarios where only scRNA-eq or scATAC-seq were downsampled can be found in Supplementary Figure S4. (B) Performance of Bridge integration under different downsampling scenarios. The error band shows the 95% confidence interval.

If we downsampled cells in the RNA or ATAC data separately, similar patterns can be observed (Supplementary Figure S4). Still, Bridge integration and Seurat v3 had the highest overall accuracy and F1 (macro), and scJoint had the highest weighted accuracy. For Bridge integration, we found that at the same downsampling rate, downsampling all cells resulted in the worst performance, followed by downsampling ATAC cells, multimodal cells and RNA cells only (Figure 5B). Therefore, Bridge integration was the most sensitive to the sequencing depth of ATAC cells and least sensitive to the sequencing depth of RNA cells of the BMMC data. For other methods, they were more sensitive to the sequencing depth of ATAC cells than that of the RNA cells as well (Supplementary Figure S4).

### 2.5 Performance When There Exist ATAC-Specific Cell Types

The last set of experiments was designed to investigate the performance of methods when there were ATAC-specific cell types by manually removing some cell types in the reference scRNA-seq data. For overall accuracy and F1 (macro), Bridge integration achieved the highest scores followed by scJoint and Seurat v3 with close performances, and Conos was still the worst performer (Supplementary Figure S5A). The overall accuracy didn’t change much as the number of removed cell types in RNA increased, while F1 (macro) decreased linearly when the number of removed cell types increased. For weighted accuracy, scJoint was the best method followed by Bridge integration and Seurat v3. For Conos, its performance became worse when Seurat was used for data processing (Supplementary Figure S5B). For Bridge integration, we found that the values of overall accuracy, weighted accuracy and F1 (macro) were smaller when removing cell types in the RNA data compared to removing cell types in the multimodal data given the same number of removed cell types (Supplementary Figure S5C).

Since after removing cell types in scRNA-seq, there existed ATAC-specific cell types, we also calculated the metrics designed for assessing performance of methods on these cell types. As shown in the last two plots in Supplementary Figure S5A, scJoint had the highest weighted accuracy followed by Bridge integration and Conos with close performances; while scGCN was the best performer in terms of F1 (entropy and enrichment) and scJoint performed worst. Therefore, scJoint tended to classify ATAC-specific cell types to their similar cell types in the reference data.

## 3 Discussion

We performed a comprehensive benchmarking study on five automated scATAC-seq label annotations methods across five different tissues using both unimodal and multimodal single-cell data. By conducting experiments on the well-annotated BMMC data, we also studied the performance across different cell numbers, mislabeling proportions, sequencing depths and number of unique cell types. We designed three overall metrics and two metrics for ATAC-specific cell types to evaluate the prediction accuracy. In addition, we assessed the running time and memory usage of each method.

Through the designed experiments on BMMC, we found that lower number of RNA cells, higher mislabeling proportions, and lower sequencing depth could lead to worse performance of all methods. When changing the number of RNA cells, we found that all methods were not sensitive to the data scale when the cell number was larger than 3k. When changing the mislabeling proportion, most methods had a significant decrease in overall accuracy and F1 (macro) only after the mislabeling proportion reached 50%. Bridge integration was able to maintain accuracy at a high level even when the mislabeling proportion was 70%. In contrast, all methods were sensitive to lower sequencing depth. Across all the experimental scenarios, we found Bridge integration was consistently the best performer in terms of overall accuracy and F1 (macro), and the second-best performer in terms of weighted accuracy. scJoint was found to always achieve the highest weighted accuracy across all experiments, suggesting it did a good job in relating similar cell types. In contrast, Conos performed the worst regardless of the processing pipeline used (Seurat or Pagoda2). Additionally, for Bridge integration, we found that the sequencing depth of scATAC-seq and multimodal data played a more important role than the sequencing depth of scRNA-seq. This might be because scATAC-seq is known to be sparser than scRNA-seq due to the limitation of current sequencing technologies (Minnoye et al., 2021).

By benchmarking across different tissues, we found that all methods had better performance than KNN and random classifiers when considering all cells. On human PBMC and BMMC where all data were measured by 10x and were published no earlier than 2019, Bridge was the leading method. However, for mouse lung and mouse brain, scJoint was the best performer. Note that the sequencing depth of SHARE-seq mouse lung data was too low so that we were not able to assess the performance of Bridge integration on mouse lung (Ma et al., 2020). For mouse brain, when applying Bridge integration, we had to remap the original fastq data of unimodal ATAC data to mm10 because there was inconsistency between the reference genome used for the provided unimodal ATAC (mm9) and multimodal ATAC data (both SHARE-seq and SNARE-seq used mm10). After remapping, we found the sequencing depth of the unimodal ATAC data was extremely low, with median count sum per cell being 78 (mapped to the peak set of SNARE-seq) and 94 (mapped to the peak set of SHARE-seq). While for other tissues, there were usually thousands of counts per cell (Supplementary Table S3). Such high sparsity might cause the poor performance of Bridge integration on mouse brain, which was consistent with the finding in BMMC experiments of changing sequencing depth. For mouse kidney, Bridge integration performed relatively badly but the difference between it and other methods was not significant, and the bad performance might also result from the low sequencing depth of multimodal RNA data (Supplementary Table S3).

For performance on ATAC-specific cell types, we found scJoint consistently had the highest weighted accuracy but the lowest F1 (entropy and enrichment), suggesting that it tended to classify unique cell types to existing cell types that were the most similar to them. On the contrary, scGCN was the best method in terms of F1 (entropy and enrichment), followed by Bridge and Seurat v3.

In terms of efficiency and scalability, scGCN was both time and memory consuming, and Conos was the most efficient algorithm. Bridge integration required additional multimodal data, so it consumed more memory than others, but its memory usage didn’t increase sharply when the data scale increased because it utilized dictionary learning and only performed heavy computation on a subset of data (Hao et al., 2022).

Our study had some limitations. First, the conclusions are tissue and technology specific. Second, the granularity of cell types was coarse for most tissues, like the three mouse tissues after unifying annotations across datasets. The performance of methods might change if finer cell annotations were provided.

Based on the findings in our benchmarking study, we have the following recommendations. If all data are from 10x and multimodal data from the same tissue are available, Bridge integration is likely the best method for label transfer; otherwise, scJoint is the to-go method. For scJoint, the caveat is that it tends to misclassify ATAC-specific cell types to the biologically similar cell types in RNA. If one cares about ATAC-specific cell types, a better strategy might be using scGCN or Seurat v3 and another method in two separate rounds. For scGCN or Seurat v3, manual annotations can be performed on cells that have high entropy and low enrichment.

## 4 Materials and Methods

### 4.1 Single-cell Data Preprocessing

A full list of data used in this study can be found in the Supplementary Table S1. Descriptions of preprocessing pipelines specific to each dataset are provided below. Moreover, to facilitate the evaluation of label prediction performance, we manually unified the naming conventions of cell labels provided in the scRNA-seq and scATAC-seq (Supplementary Table S2). Details for data preprocessing can be found in our GitHub repository.

#### Human BMMC

This is so far the largest single-cell multimodal RNA and ATAC dataset with well-annotated labels and hierarchical batch structures. To mimic the case where scRNA-seq, scATAC-seq and multimodal data were measured separately, we manually separated all batches to three groups without any overlaps. Specifically, batches s1d2, s1d3, s3d3, s4d9, and s3d10 were used as scRNA-seq (26,450 cells), s2d4, s2d5, s3d7, and s4d8 were used as scATAC-seq (22,653 cells), and s1d1, s2d1, s4d1 were used as multimodal data (18,467 cells). Since the raw gene activity matrix was not provided, the gene activity matrix for cells assigned to the scATAC-seq group was obtained using Signac (Stuart et al., 2021).

#### Human PBMC

The reference genomes used for scATAC-seq (hg19) and 10x multiome ATAC-seq (hg38) were different and only the latter had public raw sequence data in fastq formats. We remapped the 10x multiome data using cellranger-arc to get the peak count matrix and fragment files. Since Bridge integration requires that the peak sets of count matrices in scATAC-seq and multimodal ATAC data are the same, we requantified the abundance of scATAC-seq peaks on the multimodal peak set using the FeatureMatrix function in Signac. For the gene activity matrix, we used Signac to do the calculation.

#### Mouse kidney

To unify the feature set as required by Bridge integration, we requantified the scATAC-seq peaks on the multimodal peak set as what we did for human PBMC data. In addition, since the gene activity matrix for mouse kidney scATAC-seq was not provided, we calculated it using the GeneActivity function in Signac.

#### Mouse brain

The reference genome used for scATAC-seq (mm9) was different from that used for ATAC in the two brain multimodal data (mm10). To correct the inconsistency, we used the provided bam files of scATAC-seq data to map it to mm10 in three steps. First, samtools was used to convert bam to fastq files. Second, fastq files were mapped to mm10 to get new bam files using bowtie2 and samtools sequentially. Last, sinto was used to get fragment files from bam files. After getting fragment files, Signac was used to obtain the count matrix using the peak set in the multimodal ATAC data (SNARE-seq and SHARE-seq separately) and the fragment files. For the scATAC-seq gene activity matrix, we used the provided one.

#### Mouse lung

We did not find an appropriate multimodal data for mouse lung, so the data for this tissue were only used to benchmark methods that do not require multimodal data (only Bridge integration requires). For the gene activity matrix, we used the one provided by the original paper.

### 4.2 Description and Implementation of Methods

#### Conos

Conos is designed as a graph-based batch effect removal method. The joint graph embedding using nearest neighbors and Pearson correlation is constructed as the first step to connect all cells. Then, the label transfer from reference data to query data can be implemented by information propagation between graph vertices through an iterative diffusion process.

#### Seurat v3

Seurat first identifies a set of anchors between the reference and the query data through canonical correlation analysis (CCA) and mutual nearest neighbors (MNNs). Then, a weight matrix is constructed to quantify the distance between each query cell and anchor cell in the query data by a Gaussian kernel. Last, the prediction score of any cell in the query data is calculated as a weighted average of labels of anchor cells in the reference data.

#### scGCN

The first step of scGCN is to build a hybrid graph of all cells using MNNs approach and CCA. Based on the constructed graph, a semi-supervised graph convolutional neural network is trained to embed cells from both reference and query data on the same latent space and predict cell type labels for cells in the query data.

#### scJoint

Like scGCN, a semi-supervised neural network with cross entropy loss is trained to jointly embed cells from both scRNA-seq and scATAC-seq. Different from scGCN that directly utilizes the trained network to predict probability vectors through Softmax layers, scJoint performs label transfer by training an additional kNN classifier in the embedding space.

#### Bridge integration

This method utilizes multimodal data as a bridge to transfer labels from scRNA-seq to scATAC-seq. The multimodal dataset is treated as a dictionary and each cell is an atom, on which dictionary representations of both unimodal scRNA-seq and scATAC-seq are constructed. After dimensionality reduction of multimodal cells via Laplacian Eigendecompostions, unimodal cells can be embedded on the same space by the dictionary representations. Then, the final label transfer can be achieved by any single-cell integration techniques and Bridge integration chooses mnnCorrect.

For Conos, Seurat v3, scGCN and scJoint, the raw count matrix of scRNA-seq and gene activity score matrix of scATAC-seq were provided as inputs. In addition, the raw count matrix of scATAC-seq was provided for Seurat v3 to perform dimension reduction. For Bridge integration, since the information transfer was realized by using the multimodal data as a bridge, the gene activity matrix was not needed. Instead, we provided raw count matrices of scRNA-seq, scATAC-seq (mapped to the same peak set of multimodal ATAC data) and multimodal data for Bridge integration. The implementation of each method followed the instructions on their websites. Details can be found in the scripts on our GitHub repository and package versions can be found in Supplementary Table S4.

### 4.3 Benchmarking Design

To investigate the model performance across different cell numbers, mislabeling proportions, sequencing depths and number of unique cell types, we designed the following set of experiments based on the human BMMC multimodal data. For each specific setting, 20 replicates were generated using unique random seeds.

#### Change data scale

This was separated into three sub-experimental designs. (1) Change the cell numbers in scRNA-seq (reference) while keeping the scATAC-seq and the multimodal cell numbers (for Bridge integration) as 10k. The chosen numbers were 0.2k, 0.6k, 1k, 3k, 5k, and 10k. (2) Change the cell numbers in the multimodal data while keeping the scRNA-seq and scATAC-seq cell numbers as 10k. The chosen numbers were 0.2k, 0.6k, 1k, 3k, 5k, and 10k. This setting was used for Bridge integration specifically.

#### Change mislabeling proportion

The mislabeling proportions for scRNA-seq cells were chosen as 10%, 30%, 50%, 70%, and 90%. Mislabeled cells were randomly selected and assigned wrong labels based on the background compositions of other cell labels.

#### Change sequencing depth

The sequencing depths were manually changed by downsampling reads to 10%, 30%, 50%, 70%, and 90% of the original number of reads using R package DropletUtils (Griffiths et al., 2018; Lun et al., 2019). We set four different scenarios under this experiment, which are changing sequencing depth in (1) all cells, (2) RNA cells, (3) ATAC cells, and (4) multiome cells (for Bridge integration).

#### Change the number of unique cell types

We randomly removed 2, 4 or 6 selected cell types in the scRNA-seq data. Candidate cell types were those whose cell numbers were between 200 and 1,000.

### 4.4 Evaluation Metrics

#### Accuracy

After getting the predicted probability matrix across all cells in scATAC-seq, the cell type that had the highest predicted probability was assigned to each cell as the predicted label. Then the overall accuracy was calculated using the predicted labels and true labels.

#### Weighted accuracy

To account for the prediction uncertainty and similarity across cell types. We proposed a weighted accuracy (WACC) by taking the average of the predicted probability vector weighted by cell type similarities.

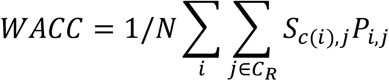

In the equation above, *P* is the predicted probability matrix with each row as a cell in scATAC-seq and each column as a cell type observed in scRNA-seq reference data. *C*_*R*_ is the set of all cell types in scRNA-seq and *N* is the total number of scATAC-seq cells. *S* is a cross-modality cell type similarity matrix with each row as a cell type in scATAC-seq and each column as a cell type in scRNA-seq and *c*(*i*) is a function mapping cell *i* to its true cell type label.

The similarity matrix was calculated in three steps. First, partition-based graph abstraction (PAGA) (Wolf et al., 2019) was performed on the normalized count matrix of scRNA-seq and gene activity matrix of scATAC-seq separately. Then, the within-modality similarity matrix was calculated based on the Euclidean distance of each pair of cell types using the PAGA positions. Specifically, we applied exp to the negative of the calculated distance matrix. Last, we calculated the cross-modality similarity matrix using the two within-modality matrices by considering three scenarios. If two cell types existed in both modalities, their similarity was calculated as the average of two within-modality similarities. If one cell type is modality-specific, its similarity with any common cell type would be the similarity calculated using the modality that contained the two cell types. If a cell type *l* only existed in scATAC-seq and the other cell type *k* was only observed in scRNA-seq, their similarity was calculated as

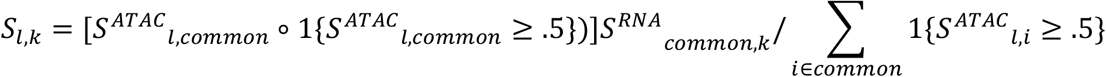

where *S*^*ATAC*^ and *S*^*RNA*^ are within-modality similarity matrix for ATAC and RNA, respectively and *common* is the set of all common cell types. The first product is Hadamard product which is element wise and the second product is matrix multiplication.

#### Precision, recall and F1 score

Precision is defined as true positive (TP) over the summation of TP and false positive (FP) and recall is defined as TP over the summation of TP and false negative (FN). F1 score is the harmonic mean of precision and recall,

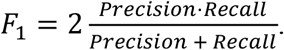

Since this is a multi-class classification problem, we need to specify whether we want macro or micro level metrics. It’s easy to show that overall accuracy is equivalent to micro precision, recall and F1 score under the multi-class scenario. Therefore, we calculated macro level precision and recall in this study, which is the average of precisions and recalls obtained for each class. Then, macro F1 score is calculated based on macro precision and recall.

#### Entropy and enrichment

To evaluate the performance of methods on cell types unique to scATAC-seq data, we borrowed the two metrics proposed in scGCN which are scaled entropy and enrichment (Song et al., 2021). Scaled entropy is defined as

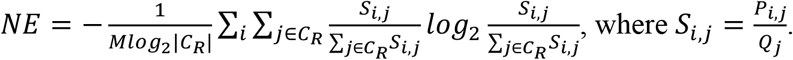

*P*_*i,j*_ is the predicted probability for cell *i* with unique cell type label in scATAC-seq and cell type *j*, and *Q*_*j*_ is the proportion of cell type *j* in scRNA-seq as the background probability. *C*_*R*_ is the set of all cell types in scRNA-seq and *M* is the total number of scATAC-seq cells with unique cell labels. The final score is normalized by *log*_2_|*C*_*R*_| to make it in the range of [0, 1]. Another metric is enrichment score,

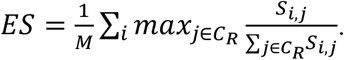

The enrichment score is also bounded within 0 and 1. For cell types only observed in scATAC-seq, an ideal method should deliver high normalized entropy and low enrichment score. Therefore, we also calculated an F1 score to combine these two

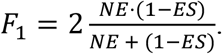

#### Running time and memory

All methods were run on Yale’s high performance computing clusters with one computing core. For neural network methods scGCN and scJoint, they were run using GPUs; and for the rest methods, they were run using CPUs. The CPU of our device is Intel ® Xeon ® Gold 6240, 2.6 GHz, and the GPU is NVIDIA RTX 3090 with 25 GB RAM. When evaluating running time, we did not count the time used for data preprocessing (e.g. remap to alternative reference genome, requantify scATAC-seq peaks, and calculate gene activity matrix) because the needed steps for different tissues were different. For memory assessment, we used the recorded peak memory usage of each method.

## 6 Data Availability Statement

All the single-cell data used in this manuscript are publicly available. Detailed information of each data and their downloadable links can be found in Supplementary Table S1. The related scripts for reproducing results in this manuscript are available on GitHub at https://github.com/AprilYuge/ATAC-annotation-benchmark.

## 7 Author Contributions

YW collected data, performed label unification and similarity matrix calculation, designed the benchmarking pipeline and evaluation metrics, assisted in preparing scripts for running each method, evaluated the model performance, and wrote the manuscript. XS wrote scripts for running each method, performed data processing and gene activity calculation, assisted in model evaluation, and provided feedback to the manuscript. HZ supervised the entire project, contributed to the design of the benchmarking pipeline and evaluation metrics, revised the manuscript critically for important intellectual content and provided approval for the publication of this manuscript.

## 9 Supplementary Material

The Supplementary Material for this article can be found online at the bioRxiv website.

