## Supplementary Figures for "Benchmarking Automated Cell Type Annotation Tools for Single-cell ATAC-seq Data"

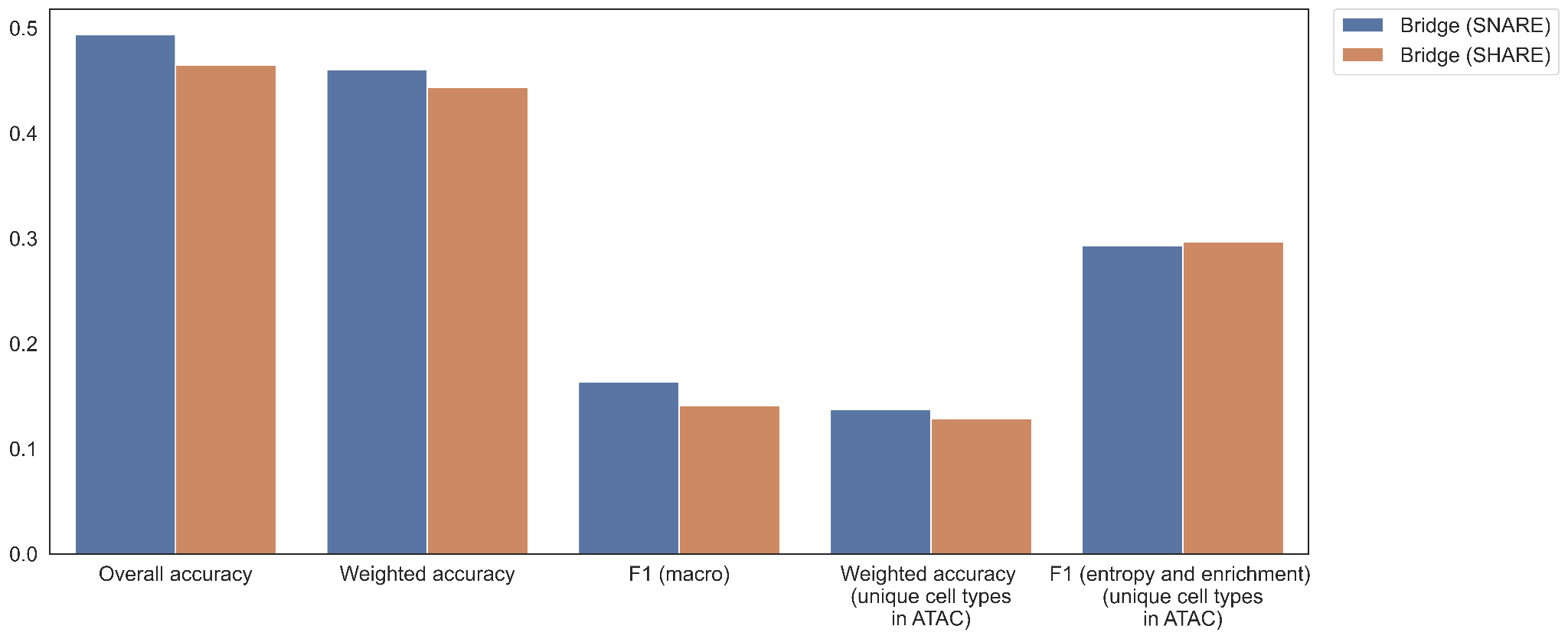

**Supplementary Figure 1.** The performance of Bridge integration on the mouse brain data with SNARE-seq or SHARE-seq data as the ‘bridge’.

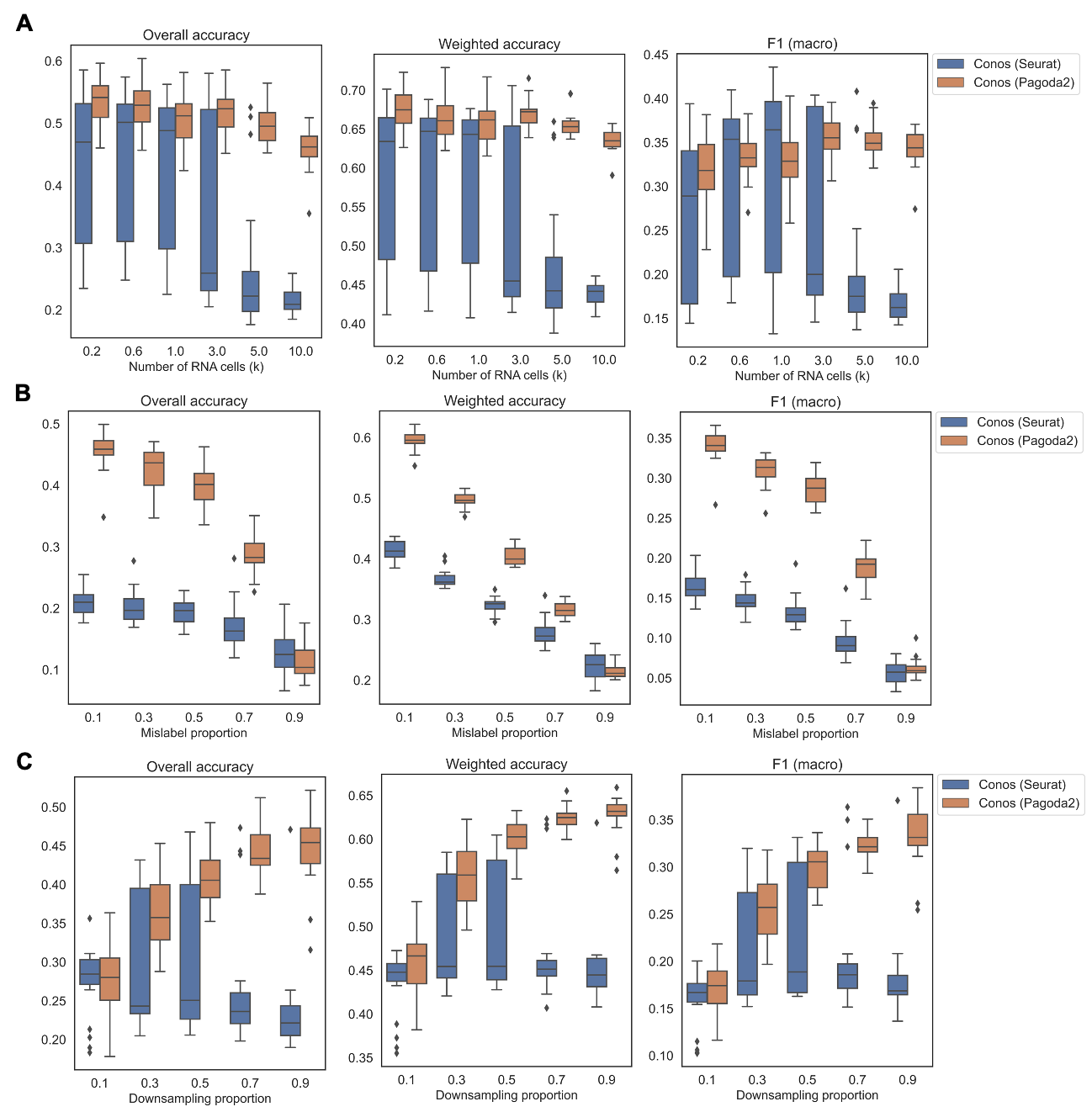

**Supplementary Figure 2.** Comparison of Conos combined with data processing pipeline by Seurat or Pagado2 across different BMMC experiment settings presented in the main text. Central lines represent medians, boxes represent the interquartile range (IQR), and the upper/lower whisker represents the largest/smallest value no further than 1.5 × IQR.

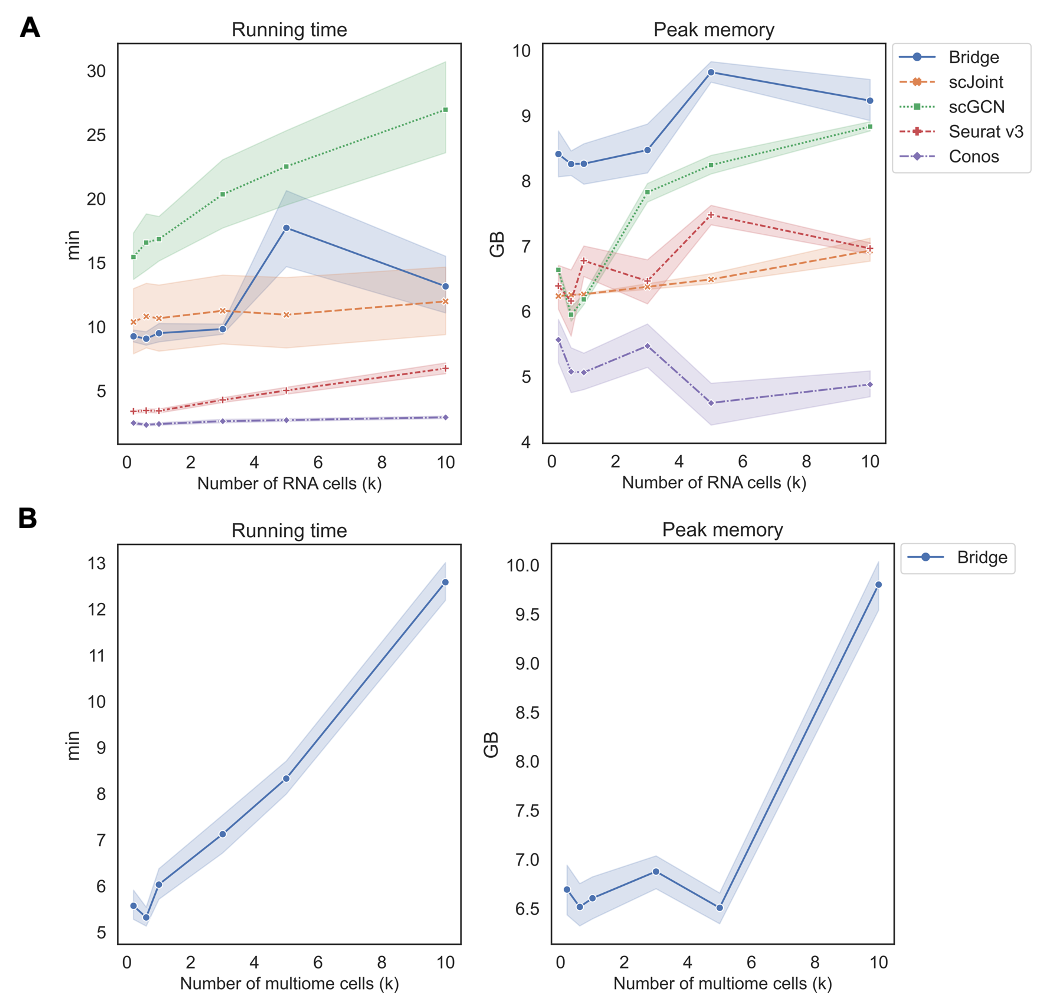

**Supplementary Figure 3.** Running time and peak memory usage of methods when changing number of RNA cells (A) and when changing number of multimodal cells (for Bridge only) (B). The error band shows the 95% confidence interval.

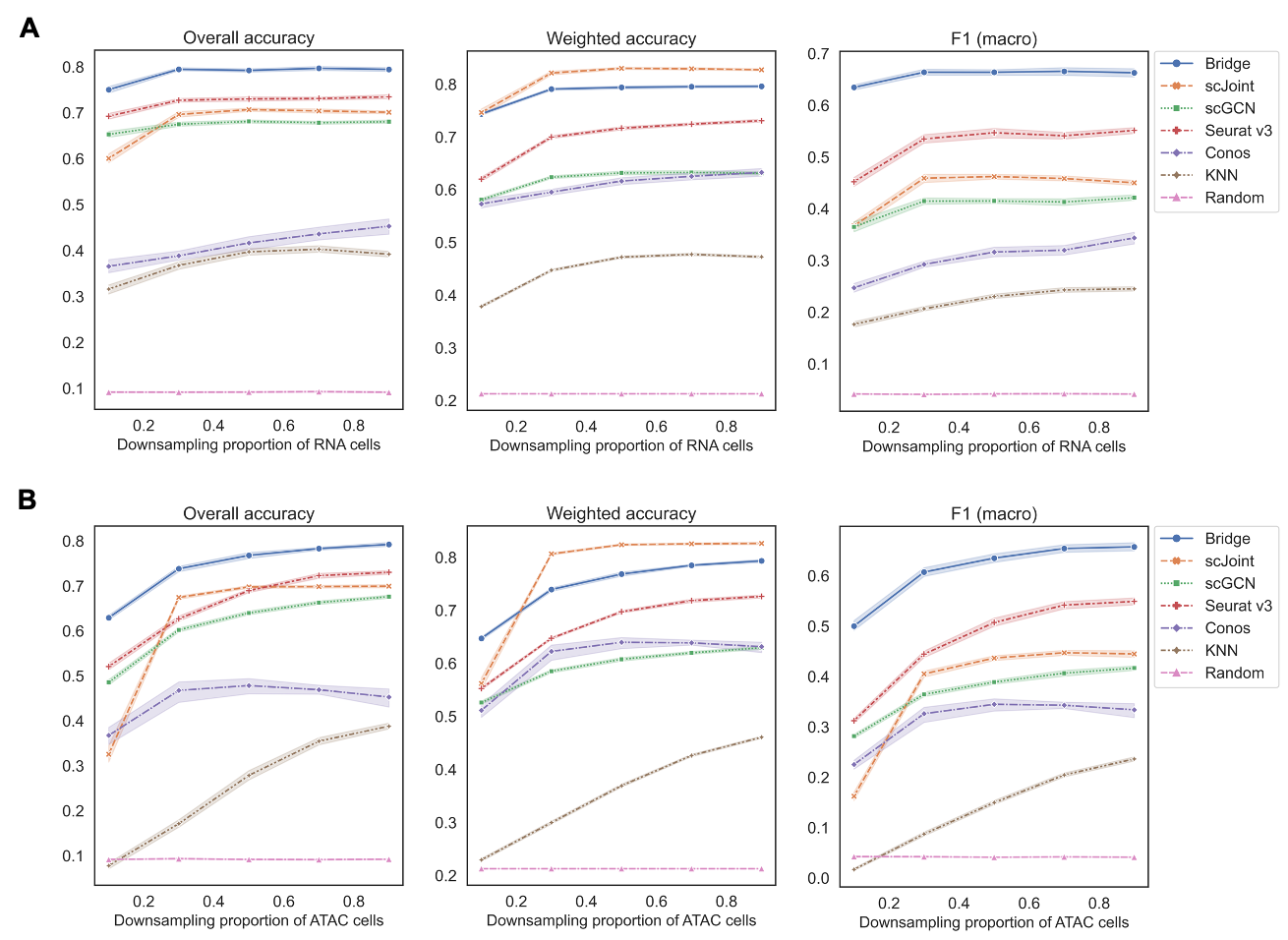

**Supplementary Figure 4.** Performance of methods on different downsampling proportions of scRNA-seq (A) and scATAC-seq (B) only, respectively. The error band shows the 95% confidence interval.

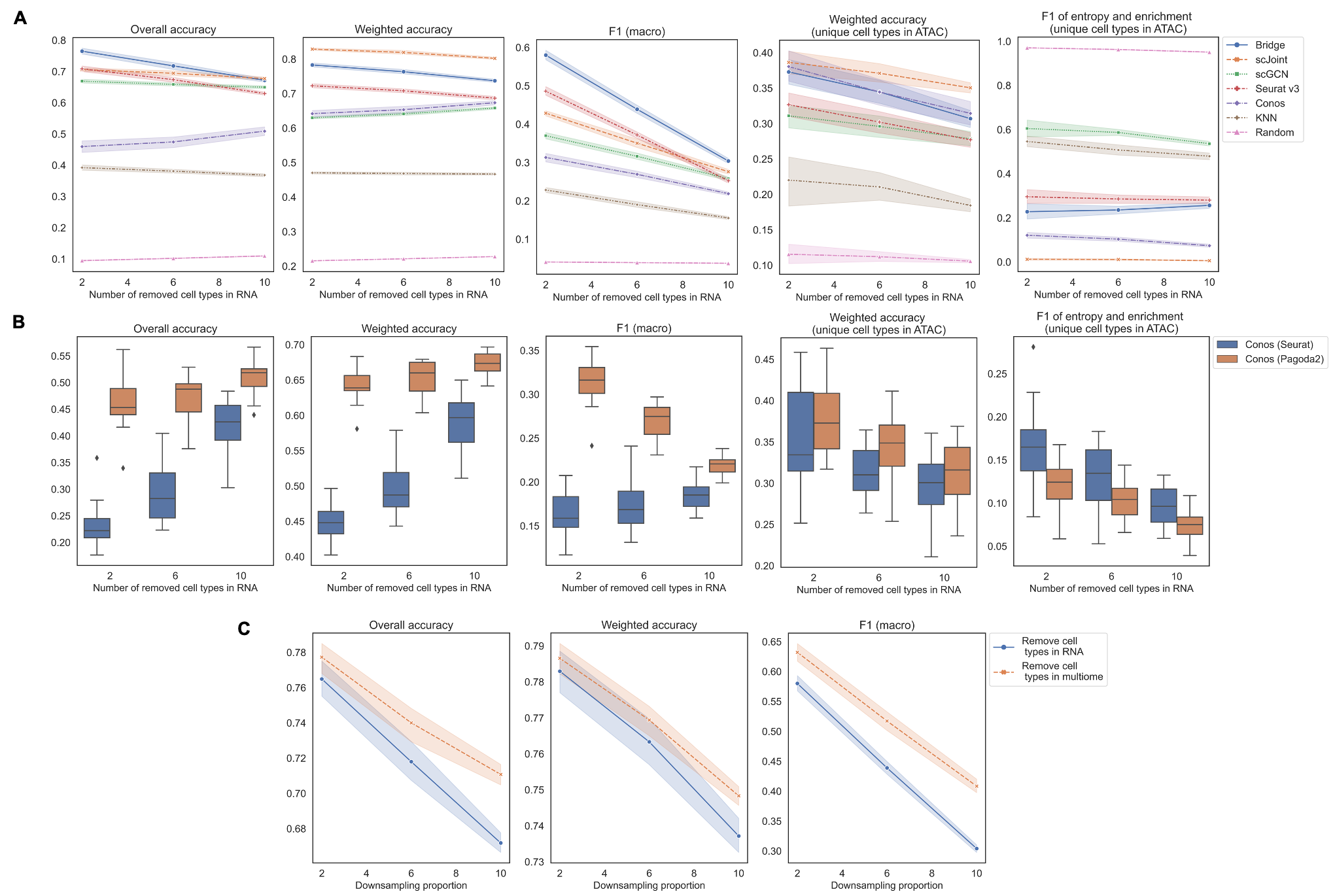

**Supplementary Figure 5.** Performance of methods on different numbers of removed cell types. (A) Remove cell types in labeled RNA data to make corresponding cell types in ATAC data unique. (B) Performance of Conos using either Seurat or Pagoda2 for data processing. Central lines represent medians, boxes represent the interquartile range (IQR), and the upper/lower whisker represents the largest/smallest value no further than 1.5 × IQR. (C) Performance of Bridge when removing cell types in RNA or multimodal data, respectively. The error band shows the 95% confidence interval.
